# Engineering the Maize Root Microbiome: A Rapid MoClo Toolkit and Identification of Potential Bacterial Chassis for studying Plant-Microbe Interactions

**DOI:** 10.1101/2023.06.05.543168

**Authors:** John van Schaik, Zidan Li, John Cheadle, Nathan Crook

## Abstract

Sustainably enhancing crop production is a necessity given the increasing demands for staple crops and their associated carbon/nitrogen inputs. Plant-associated microbiomes offer one avenue for addressing this demand; however, studying these communities and engineering them has remained a challenge due to limited genetic tools and methods. In this work, we detail the development of the Maize Root ToolKit (MRTK); a rapid Modular Cloning (MoClo) toolkit that only takes 2.5 hours to generate desired constructs (5400 potential plasmids) that replicate and express heterologous genes in *Enterobacter ludwigii* strain AA4 (*Elu*), *Pseudomonas putida* AA7 (*Ppu*), *Herbaspirillum robiniae* strain AA6 (*Hro*), *Stenotrophomonas maltophilia* strain AA1 (*Sma*) and *Brucella pituitosa* strain AA2 (*Bpi*) which comprise a model maize root synthetic community (SynCom). In addition to these genetic tools, we describe a highly efficient transformation protocol (10^7-10^9 transformants/µg of DNA) for each of these strains. Utilizing this highly efficient transformation protocol, we identified endogenous expression sequences for each strain (ES; promoter and ribosomal binding sites) via genomic promoter trapping. Overall, the MRTK is a scalable platform that expands the genetic engineering toolbox while providing a standardized, high efficiency transformation method that can be implemented across a diverse group of root commensals. These results unlock the ability to elucidate and engineer plant-microbe interactions promoting plant growth for each of the 5 bacterial strains in this study.

## INTRODUCTION

Microbiomes have diverse impacts on plant health, including nutrient acquisition (1, 2, 3), stress response (1, 4, 8), and disease resistance (1, 5, 6, 7). Determining the mechanisms behind these plant growth promoting phenotypes has been largely due to advancements in ‘omics technologies and in high-throughput phenotypic measurements. For example, 16S amplicon sequencing has greatly expanded our understanding of which microbes associate with plant hosts and which are correlated with growth promotion, while high throughput assays such as nutrient profiling (2,4), functional genomics/metagenomics (3,7), and transposon mutant libraries (5,6,8) have helped uncover the genetic bases of these properties. Many of these studies are facilitated by the use of simplified systems, such as synthetic communities or single microbes inoculated with their plant host (2,3,4,6), which greatly reduces system complexity and allows individual interactions to be studied. However, the study of plant-microbe interactions is still limited by a lack of genetic tools and methods for most members of the plant microbiome (9,10).

To bridge this gap, genetic toolkits have been developed with a variety of applications for a variety of microorganisms. We can broadly classify these toolkits into two groups: basic expression and advanced applications toolkits. For model organisms (e.g. *E. coli* (11–15), *P. putida* (16–19), *Saccharomyces cerevisiae* (20–23), etc.), a plethora of toolkits in the advanced application group exist that standardize and streamline more complex operations such as genome editing, genome integration, creating gene circuits, directed evolution, etc. A subset of these are Modular Cloning (MoClo) toolkits that utilize the golden gate cloning technique, such as the Yeast Toolkit (YTK) (20), EcoFlex (11), CIDAR (15), etc. MoClo toolkits provide standardized parts and efficient cloning strategies that make it straightforward to combinatorially assemble parts for the expression of proteins and/or the more advanced applications listed above. Genetic toolkits are also emerging for non-model microorganisms (24–29). These are usually specific to a single species and define various regulatory elements (e.g. promoters, ribosomal binding sites (RBSs) and terminators) to enable a range of expression levels and can include some advanced applications like genome editing or gene circuit design. There have also been efforts to create genetic toolkits/collections that are generalizable to larger groups of organisms like gram-negative bacteria, proteobacteria and both prokaryotes and eukaryotes (30–32). In the context of plant-associated microbes, the well-studied *Bacillus subtilis* (33–35) and *Pseudomonas* species (16–19) have genetic toolkits that allow for tuneable and inducible expression, genome engineering and protein secretion. While genetic tools are being adapted and developed in more non-model, plant-associated microbes (36, 37), single microbiome toolkits and high transformation efficiencies (a prerequisite for high-throughput genetics) remain elusive bottleneck.

To start defining genetic tools for plant microbiomes, we focused on the 7-member synthetic maize root microbiome (SynCom) developed by Niu *et al* (38). This SynCom was chosen for its simplicity, its representaton of the complex and diverse maize root microbiome, and its ability to model disease resistance and community assembly. Currently, there are 5 published studies and 1 pre-print (BioRxiv) (6, 39–43) that have utilized this SynCom, indicating that it is an emerging standard for plant microbiome research. This SynCom has enabled studies of community stability/dynamics (39 & 40), microbe-dependent heterosis in maize (41), development of protein extraction protocols for meta-proteomics in maize (42), identification of biocontrol strains against nematodes (6), and metabolic analysis of the SynCom strains (43). Genetic tools are available for two SynCom species: *Enterobacter ludwigii* strain AA4 (*Elu,* formerly *E. cloacae*) and *Pseudomonas putida* strain AA7 (*Ppu*) where genetic tools/methods developed in *E. coli* have been shown to be adaptable for Enterobacter strains (44–46) and a plethora of tools developed in other Pseudomonas putida strains that may be adapted to strain AA7 (16–19). The rest of the SynCom, *Stenotrophomonas maltophilia* strain AA1 (*Sma*), *Brucella pituitosa* strain AA2 (*Bpi,* formerly *Ochrobactrum pituitosum*), *Curtobacterium pusillum* strain AA3 (*Cpu*), *Chryseobacterium indologenes* strain AA5 (*Cin*) and *Herbaspirillum robiniae* strain AA6 (*Hro,* formerly *H. frisingense*), vary in the availability of genetic tools. Transposon mutants have been generated in *Herbaspirillum* (*51,52*) and *Stenotrophomonas* species (47,48), expression vectors have been constructed for *Brucella* species (49,50), but no tools are available for *Chryseobacterium* and *Curtobacterium* species to the author’s knowledge. Furthermore, transformation methods have yet to be established for *Chryseobacterium* species, but either electroporation and/or conjugation has been used to transform species in the same genera for the rest of the SynCom (53–58).

In the work presented here, we outline a set of genetic tools and a universal, highly efficient transformation method for 5 of the gram-negative SynCom members (*Elu*, *Ppu*, *Hro*, *Sma* and *Bpi*) to act as a foundation for future high-throughput characterization and interrogation of plant-microbe colonization & interactions in the maize root microbiome. Specifically, we developed a standardized MoClo plasmid toolkit for the strains *Elu*, *Ppu*, *Hro*, *Sma* and *Bpi* that enables rapid cloning directly into these SynCom members in about 2.5 hours. Additionally, we have developed a high-efficiency transformation method and used it as a proof-of-concept to identify expression sequences (ESs; specifically promoters and RBSs) via promoter-trapping screening, enabling tuning of protein expression in each of the strain.

## RESULTS

### The Maize Root ToolKit (MRTK)

The maize root toolkit (MRTK) is a MoClo toolkit that allows users to create their desired product plasmids (P plasmids) with the MRTK cloning plasmids (C plasmids) and transform their P plasmid into their desired organism within 2.5 hours (Figure 1A). Specifically, the MRTK consists of 45 C plasmids (Table S1) that follow the common PCR-free format of MoClo toolkits (20, 63) and that can be used to create 5400 total P plasmids. The MRTK C plasmids contain one of 5 different part types (origin of replication, selection marker, terminator, expression sequence, or fluorescent protein). All C plasmids, annotated as Clo##, contain the R6kγ origin of replication (with the exception of Clo1-Clo3 and Clo1’-3’) and are maintained in the Invitrogen *E. coli* PIR1 cell line. The R6kγ origin of replication requires the PIR gene for replication and since *Elu*, *Ppu*, *Hro*, *Sma* and *Bpi* do not contain the PIR gene, plasmids containing this origin are not maintained in these strains. This allows for the direct use of the C plasmids as starting material for cloning without the risk of them being replicated in the desired SynCom strain. Clo1-3 and Clo1’-3’ contain the Broad Host Range origin (BHR1), a temperature sensitive origin (SC101+RepA protein) or the ColE1 origin with the chloramphenicol (Clo1-3) or ampicillin (Clo1’-3’) selective markers. These two sets of plasmids were generated to allow users to have a selection marker for their P plasmid that is different from the origin-containing C plasmids used for cloning. Multiple origins also allow for the creation of dual plasmid systems often seen in genome editing systems (12, 46).

**Figure 1:**
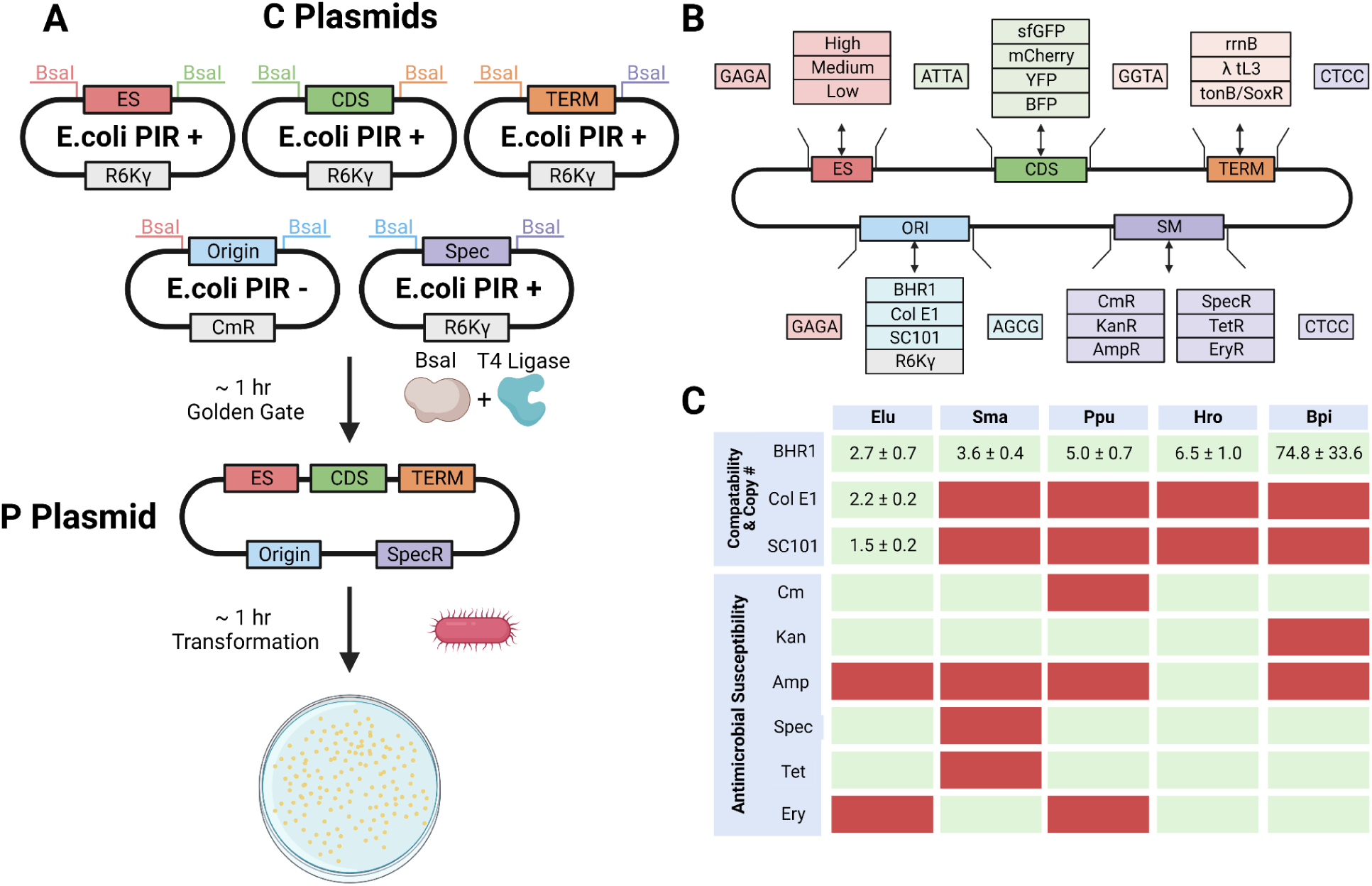
The Maize Root Tool Kit (MRTK) An overview of the MRTK cloning process, parts within the toolkit, origin compatibility, origin copy number and antimicrobial susceptibility for each strain. A) Cloning plasmids (C plasmids) are used as starting material with BsaI restriction enzyme and NEBridge MasterMix to clone the product plasmid (P plasmid). P plasmids can then be transformed into the desired SynCom strain of interest with a time of ∼2.5 hours from plasmid to plate. B) Overview of all 45 parts included in the MRTK and their 4 bp golden gate overhang sequences. C) A table showing the compatibility and copy number of the origins of the MRTK for each strain as well as their antimicrobial susceptibility. Green boxes indicate that the origin is replicated in the strain and that the strain is susceptible to the antimicrobial agent. Red boxes indicate that the origin is not replicated in the strain and that the strain will grow on that antimicrobial agent.

All C plasmids can be cloned into a P plasmid using golden gate cloning with the restriction enzyme BsaI. While standard golden gate methods work with the MRTK, we have found that using the NEBridge Master Mix enables a plasmid-to-plate time of 2.5 hours. If users wish to utilize parts that are not provided with the kit (Figure 1B), we suggest manual PCR with the appropriate golden gate overhangs. The golden gate overhangs for the MRTK were selected based on the high fidelity overhangs listed in Supplementary Table S8 from Potapov et al (64). The selective marker compatibility, copy number, and origin compatibility were determined for each of the 5 strains (Figure 1C). We found that all origins are maintained at low copy numbers in each of the strains and that all strains are sensitive to the antibiotic combinations of spectinomycin or tetracycline combined with kanamycin, chloramphenicol or erythromycin, making these combinations good selective markers for future microbiome experiments using the SynCom.

The MRTK has several features that enable further expansion. Specifically, Clo2 and Clo3 contain the SC101 and ColE1 origins, respectively, that can replicate in *Elu.* These origins are present in an inducible CRISPR-Cas toolkit that has been used for genome editing in *Enterobacter cloacae* FRM (46). While not included in this set of vectors, the MRTK is designed to enable implementation of these promoters and CDSs through PCR with proper golden gate overhangs. Furthermore, incorporation of other CRISPR tools such as the dual-plasmid, CRISPR-prime editing system developed by Tong et al could also be incorporated through similar means (12). Additionally, Clo4 contains the non-replicative origin R6Kγ plus an origin of transfer (oriT) sequence. While the non-replicative origin enables the rapid cloning protocol, the oriT sequence on Clo4 enables its use for conjugation and has been employed in genomic editing methods such as transposon-mutant libraries (6). These are important functionalities for plasmid systems as deep understanding of plant-microbe interactions can come from genetic knockouts/knock-in experiments (65,66).

### Standardized Transformation Protocols for High Efficiency Transformation

To enable more advanced experimentation such as genome editing and high-throughput screening, high efficiency DNA transformation is required. Transformation efficiency varies drastically between microbes and based on the transformation method employed, thus optimization of a DNA transformation protocol is typically developed separately for each strain (54–56). To streamline experimentation within the SynCom, we aimed to develop a standardized transformation protocol that works for each strain of the SynCom to enable high-throughput experimentation. Electroporation was chosen over other methods of DNA transformation as it has offered the highest transformation efficiency for the majority of SynCom relatives in prior work (54–56, 67). We first tested different electroporation voltages for each strain (Figure 2A).

**Figure 2:**
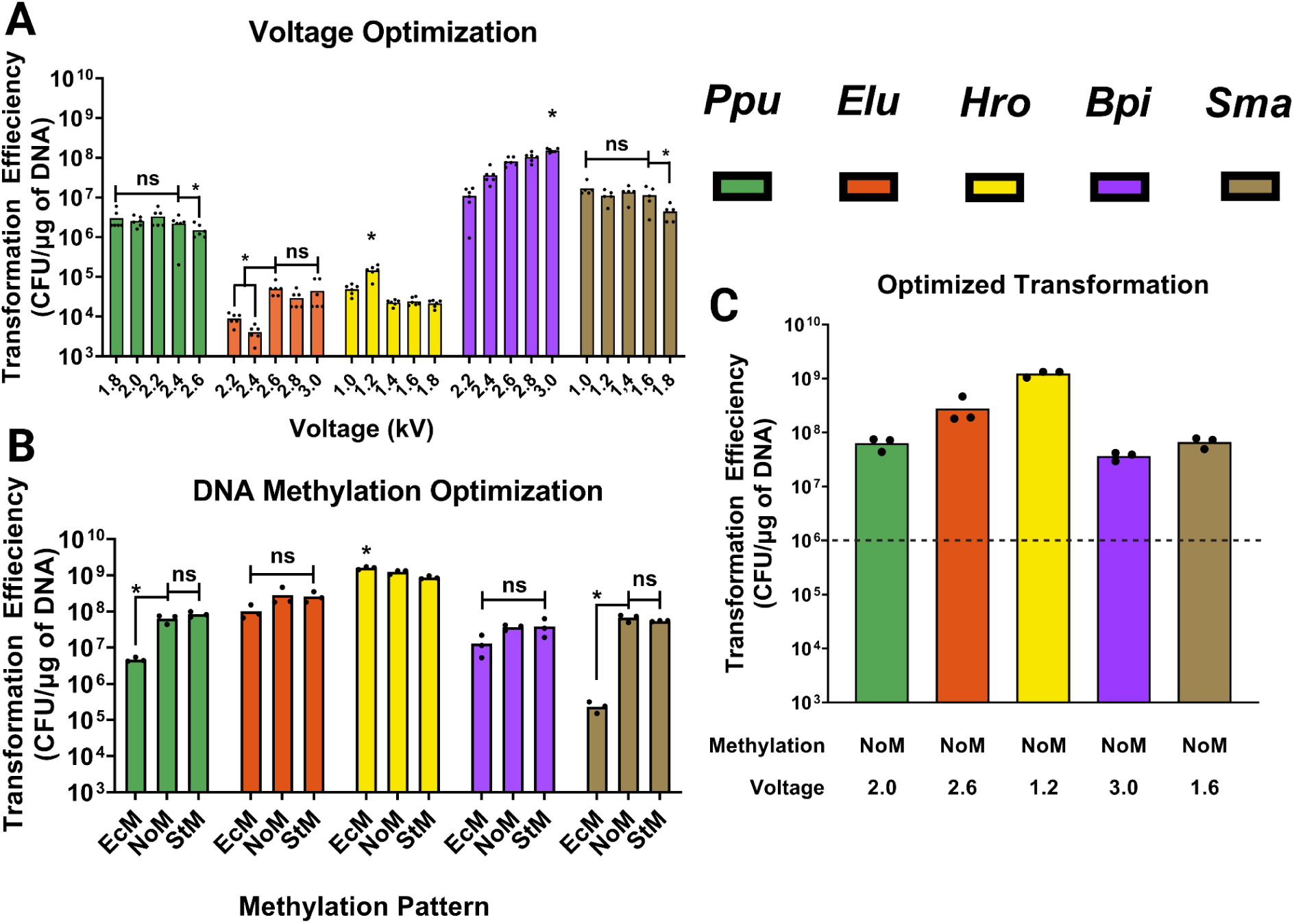
High Efficency Transformation of SynCom strains. A) Electroporation voltages for each strain. * denotes the optimal voltage or significant differences between the averages. ns denotes no significant differences between the averages. Each dot represents a technical replicate (n=5) Note: The optimal range of voltages shown for Ppu & Sma were determined from the extended range of voltages in Figures S1 & S2 B) Methylation pattern of transformed plasmids optimized for each strain. *E. coli* methylation pattern (EcM), no methylation pattern (NoM) and strain methylation pattern (StM). Each dot represents a biological replicate (n=3) C) The highest efficiency transformation protocol for each strain based on the methylation pattern of the transformed DNA and the voltage. Each dot represents a biological replicate (n=3). * denotes the optimal voltage/methylation or significant differences between the averages. ns denotes no significant differences between the averages. For 1A and 1B, optimal conditions were determined by One-way ANOVA with multiple comparison using Tukey-Kramer and HSU’s MCB in JUMP Pro17.

The strains *Bpi* & *Hro* had clear optimal voltages of 3.0 and 1.2 kV, respectively, with the rest of the strains having a larger range of voltages being optimal. Specifically, *Sma*, *Ppu* and *Elu* had a range of voltages (1.0 - 1.6 kV, 1.8-2.4 kV and 2.6 - 3.0 kV respectively) that were significantly higher from the rest of the tested voltages (Figure 2A). Additional voltages tested in *Ppu* and *Sma* are shown in Supplemental Figures S1 & S2.

We next asked whether each strain’s endogenous restriction modification (RM) systems significantly impact transformation efficiency. Briefly, RM systems are native defense systems that protect the host from foreign DNA (identified via its methylation pattern) and have been shown to have significant effects on transformation efficiency (59–62). The impact of DNA methylation on transformation efficiency was tested using DNA extracted from NEB’s *E. coli* DH5**α** strain (containing the Dam and Dcm methyltransferases, EcM), *E. coli* 135 (an *E. coli* mutant without any restriction-modification systems, NoM), and the SynCom strain of interest (StM). The impact of methylation varied across each strain, with a significant decrease in efficiency occurring for EcM in *Sma* and *Ppu* (Figure 2B). Interestingly, none of the transformation efficiencies were enhanced by the endogenous methylation patterns of each strain versus NoM, despite the existence of multiple RM systems in each strain (Supplementary Table S4). Overall, the most efficiently transformable strain was *Hro* with an efficiency >10^9 CFU/µg of DNA. This rivals the efficiency of commercially-available electrocompetent *E. coli* cells which often have transformation efficiencies of 10^9-10^10 CFU/µg of DNA. Furthermore, *Hro* lacks endogenous RM systems, making it a prime candidate for shuttling DNA libraries to other strains, especially since we found the transformation efficiency of *E. coli* 135 to be relatively low (<10^3 CFU/µg). Taken together, these results provide protocols to deliver DNA into the SynCom at high efficiency, which is a prerequisite for utilizing advanced genetic tools such as genome editing or variant screens.

### High-throughput Functional Screening for Endogenous Expression Sequences

Next, we leveraged the highly efficient transformation protocol to curate a library of expression sequences for each of the 5 SynCom strains. To do this, we used a promoter trapping method in which randomly-generated genomic fragments from each SynCom strain were ligated upstream of sfGFP, generating a library which was directly transformed without intermediate passage through *E. coli*. Transformants were sorted using fluorescence-activated cell sorting (FACS) (Figure 3A) and colonies appearing green when plated were isolated and sequenced (Figure 3B). Repair of the resulting sequences was often necessary as the ligated fragments would sometimes contain part of the downstream gene they transcribed. In these instances, the sequences were “repaired” by deleting the downstream gene fragment, and the fluorescence intensities of the repaired expression sequences were re-measured using flow cytometry.

**Figure 3:**
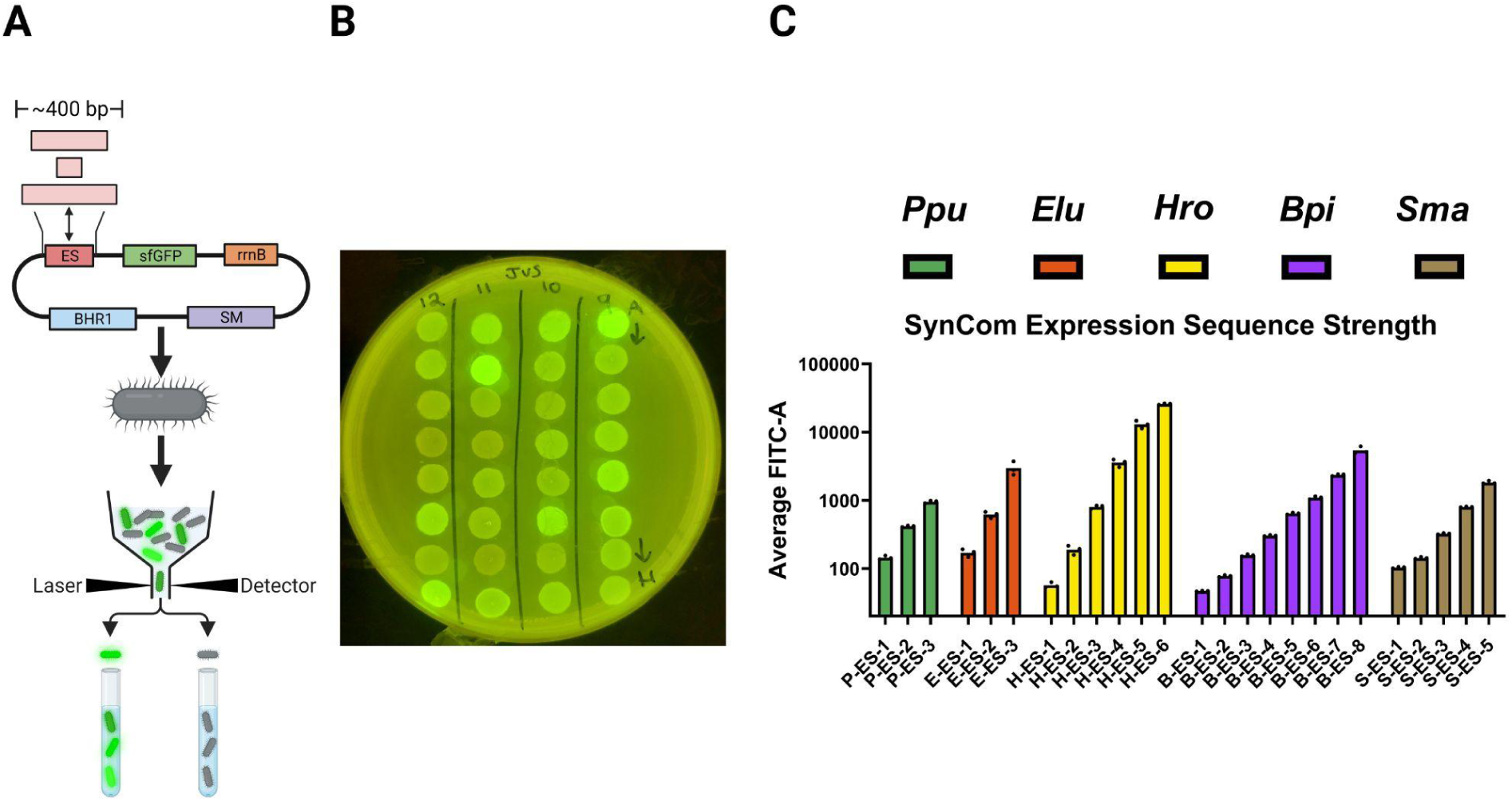
High-Throughput Expression Sequence Determination. A) Overview of the promoter trapping method coupled with FACS used to screen for variable expression sequences within the 5 SynCom strains. Genomic DNA fragments from 200-600 bp in length were ligated into a vector containing sfGFP and transformed into the strain of interest. Transformants were then sorted for varying degrees of fluorescence by FACS. B) Image of 24 *Hro* isolates after FACS sorting C) The average FITC-A values of sfGFP being expressed with each ES in their respective strains. All ESs were run in biological triplicates in their respective strain.

Collectively, we found 3-8 expression sequences for each of the 5 transformable strains that covered a 5-300 fold range of expression levels, without the need to obtain transcriptomic or proteomic data (Figure 3B). The most common downstream genes for the collection of ESs across the SynCom strains were ribosomal proteins, transporter proteins and proteins with no known function (Supplemental Table 5). The transformation protocols we identified enabled us to obtain complete coverage of each strain’s genome in these screens, with over 100-fold coverage of *Bpi* (Table 1). It is important to note that direct transformation of the cloning reactions into *Sma* or *Bpi* yielded very poor results, with 10% and 40% coverage of their genomes, respectively (Supplemental Figure 1). To overcome this, we transformed the cloning reactions directly into *Hro* and then transformed the extracted DNA to the respective strain for sorting. This boosted the coverages of each genome over 200-fold; enabling us to screen for more expression sequences and providing more evidence that *Hro* is very useful for shuttling experiments (Figure 3C).

**Table 1:** Expression Sequence Library Statistics. Library coverage is based on the total transformants, the average bp of the inserts and the genome of the strain. Colonies sequenced represents the colonies selected from the sorting plates that were sequenced.

|  | Genome Size<br>(Mbp) | Library<br>Coverage | Number of sfGFP<br>Cells sorted | Colonies<br>Sequenced | Repaired<br>ES |
| --- | --- | --- | --- | --- | --- |
| *Elu* | 4.80 | 6.08x | 7,076 | 20 | 3 |
| *Sma* | 4.66 | 46.83x | 2,011 | 17 | 5 |
| *Ppu* | 6.14 | 4.65x | 1,982 | 20 | 3 |
| *Hro* | 5.45 | 37.68x | 4,326 | 7 | 6 |
| *Bpi* | 5.47 | 107.55x | 910 | 15 | 8 |

## DISCUSSION/CONCLUSION

This report presents a rapid, modular cloning system and highly efficient transformation methods for 5 bacterial strains within the maize root microbiome. It provides an array of fluorescent proteins and the ability to tune expression levels over a wide range for each strain. This modular format allows users to integrate any sequence of interest not already within the toolkit with ease. Utilization of the high transformation efficiency protocol allowed us to find a range of expression sequences for each strain and demonstrated its potential for future use in high throughput experimentation such as mutant-library generation, functional metagenomics/genomic screening, genome editing, directed evolution, etc.

While the scope of this work was focused on providing a standardized platform for engineering multiple strains in the SynCom, its applicability may not end with the five strains studied here. Proteobacteria are one of the most dominant bacterial phyla of the Maize root microbiome (38,68,69). Since these five strains represent a diverse set of proteobacteria, it is reasonable to hypothesize that the method and parts toolkit would also efficiently transform and function within other soil bacteria. In fact, most members of microbial communities have been shown to be transformable by either electroporation or conjugation protocols (48, 54, 55, 70, 71); even if the conditions require optimization for high efficiency in some members. Further optimization of other transformation variables not addressed here (final pellet optical density, time of recovery, growth media, etc.) and incorporation of conjugation would greatly aid in enhancing and potentially enabling transformation of other members of the maize microbiome not studied here. Additionally, further expansion of the toolkit to include parts such as inducible promoters and other origins of replication would enable more complex engineering study of microbe-plant interactions within the maize SynCom as well as other rhizosphere bacteria.

RM-systems often prevent high-throughput screening within non-model organisms more generally, which we observed in our methylation experiments and when constructing ES libraries for *Sma*. In this study, we were able to overcome difficulties by shuttling the libraries through *Hro*. The high efficiency of transformation into *Hro,* coupled with non-digesting RM systems (Supplemental Table S4), makes this strain an ideal candidate for shuttling libraries to non-model bacteria. Most functional screening studies utilize commercially-available *E. coli* to generate and shuttle the libraries they wish to screen due to its high transformation efficiency (5, 72 & 73). While the total size of the libraries generated using *E. coli* is large, the total number of transformants screened may drop drastically when shuttled to a screening strain that is sensitive to *E. coli*’s methylation pattern. To get around this, the *E. coli* strains have been used as the screening strain. This is problematic for screening as the context for expressing these libraries is important. Since *E. coli* are often not the native host for genes/gene clusters found in these libraries, they may not be expressed and therefore lost during the screening process (74). For screens of plant microbiome metagenomes specifically, *Hro* could address these concerns as it is a native member of the maize root microbiome, does not have active RM systems that would inhibit transformation into other strains, and has the potential to express genes not actively expressed in competent *E. coli* strains. Furthermore, *Herbaspirillum* strains themself have been shown to be an important member of microbiomes. Specifically, *Herbaspirillum* strains promote plant growth and fix nitrogen for not only maize roots but other plant hosts (75–80). Taken together, *Hro* has great potential for engineering the microbiome and acting as a potential chassis for understanding plant-microbe interactions.

The ESs defined in this study showed stable expression in each of the strains under the culture conditions we employed. While many of the ESs are upstream of housekeeping genes (ribosomal proteins), there are a significant number that are not, and their strength may be context-dependent. For example, the expression sequence giving the highest fluorescence values we observed was H-ES6 from *Hro*’s genome. This ES lies ∼1.3 kb upstream of a predicted gene of unknown function. Furthermore, this gene was located in a ∼35.5 kb cluster of genes with unknown functions in *Hro*’s genome. While this result is intriguing, more work is needed to understand the nature of this gene cluster, and the activity of all ESs should be further validated under a variety of culture conditions and, perhaps more importantly, *in planta*.

Collectively, we envision the MRTK as greatly facilitating engineering of the maize microbiome and experimentally querying microbe-microbe and plant-microbe interactions. This toolkit expands the taxonomic range of microbe engineering within the plant-microbiome space where tools and methods are currently sparse, and provides a basis for further addition of advanced genetic engineering and control methods.

## METHODS

### Bacterial strains, media, and growth conditions

The SynCom strains used in this study were *Stenotrophomonas maltophilia* AA1 (*Sma*), *Brucella pituitosa* AA2 (*Bpi*), *Enterobacter ludwiggi* AA4 (*Elu*), *Herbaspirillum robineae* AA6 (*Hro*), and *Pseudomonas putida* AA7 (*Ppu*), all of which were obtained as a generous gift from Dr. Manuel Kleiner. *Escherichia coli* strains used were DH5α (NEB cat C2987H), *E. coli* PIR1 (Invitrogen cat C101010) and *E. coli* 135 (kind gift of Dr. Chase Beisel). All *E. coli* strains were grown in lysogeny broth media (LB) from BD scientific (BD 240210) and incubated at 37 °C and 250 rpm. Antibiotic-containing media were made with the following concentrations: chloramphenicol (Cm; 34 mg/mL), kanamycin (Kan; 50 mg/mL), ampicillin (Amp; 100 mg/mL), spectinomycin (Spec; 150 mg/mL), tetracycline (Tet; 10 mg/mL) or erythromycin (Ery; 250 mg/mL). All SynCom strains were grown with tryptic soy broth without dextrose (TSB; BD 286220) and incubated at 30 °C and 250 rpm. All plates were made according to manufacturers recommendations with 1.5% agar. The SynCom strains *Elu*, *Sma* and *Ppu* grown on TSB plates were incubated overnight at 30 °C while *Bpi* and *Hro* were incubated for 2 days at 30 °C. All SynCom strains were plated from −80 °C stocks. Transformation recovery media (SOC Media) was made by autoclaving 28 g of SOB Broth powder in 1 L of dH2O and, once cooled, 20 mL of sterilized 1 M MgSO_4_ and 20 mL of sterilized 1 M glucose were added.

### Plasmids & Primers

All primers and plasmids used for the construction of the C plasmids, transformation optimization experiments, and expression sequence experiments are listed in Supplementary Table S2 and S3 respectively of the supplementary excel file. C plasmids can be found in Supplementary Table S1. All C plasmids can be found on Addgene ######.

### Molecular cloning

Colony PCR reactions for sequence verification were performed with NEB OneTaq HotStart 2x MasterMix (M0484S) and PCR reactions for molecular cloning were performed with Q5 HotStart 2x MasterMix (M0494S). All PCR reactions were performed according to the manufacturer’s protocol. All plasmids were verified using a combination of whole-plasmid sequencing from Plasmidsaurus and Sanger sequencing from Genewiz (Azenta Life Sciences). Molecular cloning reactions were performed with either NEB Q5 mutagenesis kit (E0552S), NEB HiFi Master Mix (E2621S), or NEBridge MasterMix (M1100S) according to the manufacturer’s protocol for each kit/master mix. Molecular cloning for creating the toolkit C plasmids were performed with the NEB HiFi master mix. Molecular cloning utilizing the C plasmids were performed with NEBridge Mastermix and equimolar amounts of each C plasmid. For most of the cloning plasmids maintained in Invitrogen’s *E. coli* PIR1 cells, plasmid dimers formed and were accounted for when adding equal molar amounts of C plasmids during cloning. The length of each C plasmid is included in Supplementary Table 1. The plasmids that were used to clone the plasmids created in this study can be found in Supplementary Table S3 (20, 55, 74–78).

The empty vectors used for optimizing the transformation protocol were cloned using the NEB Q5 mutagenesis kit and Gibson cloning using NEB HiFi master mix starting from the plasmid pBBR122. Annotation of pBBR122 using plannotate (75), revealed that the Kan gene split pBBR122’s origin of replication into two parts (Supplementary Figure S4). In order to generate the empty vectors used during transformation (only the origin of replication & a single selection marker), a series of 3 cloning reactions were conducted. The first reaction removed the MobA gene sequence to generate pBBR122-ΔMobA. The second cloning reaction inserted the rrnB and T7Te terminators from pBbB1c-GFP between the origin of replication and the Cm resistance gene to generate pBBR122-DT-ΔMobA. The third cloning reaction removed the KanR gene to generate pBBR122-CmR. A set of cloning reactions (Reaction 4 in Figure S4) swapped out the CmR gene via gibson cloning with the antibiotic resistance genes for Kan, Amp, Spec, Tet and Ery. A total of eight plasmids were generated from pBBR122 with pBBR122-CmR and pBBR122-KanR used for optimizing the transformation protocol.

In order to screen the genome of each strain for functional ESs, the base vectors pBBR122-CmR-sfGFP and pBBR122-KanR-sfGFP were created from pBBR122-CmR, pBBR122-KanR and pYTK0047 using Gibson assembly using NEB HiFi Master Mix. Specifically, the promoter sequence, ribosome binding site and sfGFP coding sequence from pYTK047 were cloned in between the T7Te and rrnB terminators in pBBR122-CmR and pBBR122-KanR using Gibson assembly according to the manufacturer’s protocol.

Two sets of plasmids were created for the plasmid copy number assay. The first set of plasmids that were generated (pBBR122-CmR-sfGFP’, pBBR122-KanR-sfGFP’, pSC101-CmR-sfGFP’ and pColE1-CmR-sfGFP’) were for transformed into each strain and cloned with Gibson assembly using NEB HiFi Master Mix. The exception was pBBR122-CmR-sfGFP’ as it was generated while screening for ESs in *Elu*. The plasmid pBBR122-KanR-sfGFP’ replaced the promoter and ribosome binding site of pBBR122-KanR-sfGFP with the *Elu* ES3 from pBBR122-CmR-sfGFP’. The plasmids pSC101-CmR-sfGFP’ and pColE1-CmR-sfGFP’ replaced the BHR1 origin with the SC101 and ColE1 origins from Clo3 and Clo2 respectively. The second set of reference plasmids were generated (pZL266-Elu-DAHP-sfGFP, pZL296-Hro-rpoS-sfGFP, pZL300-Bpi-rpoN-sfGFP, pZL293-Sma-rpoD-sfGFP, pZL299-Ppu-rpoD-sfGFP) to create a standard curve for qPCR. This second set used pBBR122-KanR-sfGFP as a backbone and contained the qPCR reference genes (Supplementary Table S6) via gibson cloning using NEB HiFi MasterMix.

### Plasmid Copy Number assay

Plasmid copy number was quantified using quantitative PCR (qPCR). Each strain was transformed with either pBBR122-CmR-sfGFP’ or pBBR122-KanR-sfGFP’. *Elu* was additionally transformed with pSC101-CmR-sfGFP’ and pColE1-CmR-sfGFP’. Three individual colonies for each strain were inoculated into liquid TSB from a freshly streaked TSB plate with the appropriate antibiotic and grown at 30 °C overnight. Cultures were subinoculated to an initial OD_600_ = 0.01 in 25 mL TSB with Cm or Kan and grown to late log phase for genomic DNA extraction. *Ppu* was the only strain transformed with pBBR122-KanR-sfGFP’ and grown in TSB with Kan. Total DNA was isolated from 1 mL of late log phase cell cultures using Zymo Quick-DNA™ Miniprep Kit (D3025) according to the manufacturer’s protocol. The reference plasmids, containing sfGFP and each strain’s reference gene, were grown overnight in LB media and extracted using Zyppy Plasmid Miniprep Kit (D4020).

Since the sfGFP gene is present in all plasmids, a section of the gene was used to determine plasmid copy number. For generating the standard curve, a single copy gene in each strain’s genome was selected as the reference gene (Supplemental Table S6). Primers for each strain’s reference gene were designed using Primer-Blast with default settings for annealing temperatures of 60 °C (Supplemental Table S2). Serial dilutions of the plasmids used for standard curve generation were made with concentrations ranging from 10^4 copies/µl to 10^9 copies/µl.

Quantitative PCR was performed on a Biorad FX384 Touch Real-Time PCR Detection System. A qPCR mixture of 6 µl was prepared using SsoAdvanced Universal SYBR® Green Supermix (Cat #1725270): 0.7 µL PCR grade water, 0.15 µL of each primer (final concentration 0.375 µM), 3 µL Supermix and 2 µL template DNA. All analyses of the qPCR curves were performed using Microsoft excel.

### Optimized Electroporation Protocol

Single colonies of each strain were grown in 3 mL TSB media overnight (12-18 hours) and sub-inoculated at a 1:40 ratio into 25 mL TSB media the next day. 1 mL of each overnight culture was aliquoted for genomic DNA extraction using Zymo’s Quick-DNA Fungal/Bacterial Miniprep Kit (D6005). Extracted genomic DNA was used for 16S PCR with standard 27 forward and 1492 reverse primers to verify purity of the culture to be transformed. PCR conditions followed the standard protocol outlined for NEB OneTaq HotStart 2x Master Mix with an annealing temperature of 49 °C, a cycle extension time of 1 minute 30 seconds, and a final extension time of 2 minutes. Cultures were harvested at an OD_600_ of 0.4-0.6 by centrifugation at 5,000 xg for 5 min at 4 °C. The supernatant of harvested cells was decanted and the pellet was resuspended gently with a pipette using 25 mL of ice-cold 10% glycerol. The cells were washed by this process three times. Cells were finally resuspended with 250 μL of ice-cold 10% glycerol and normalized to an OD_600_ of 25. 50 μL of the normalized cell suspensions were electroporated with 2 μL of plasmid DNA (up to a maximum of 5 μL) at the respective strain’s optimal voltage in 1 mm gap electroporation cuvettes (USA Scientific Inc 91041050) with a Biorad MicroPulser electroporator. Electroporated samples were recovered at 30 °C and 250 rpm for 1 hour in 950 μL of SOC media. After recovery, serial dilutions of cell recovery solutions were made in a 96 well plate with total volumes of 250 μL. 10 μL of each serial dilution were then spot plated onto a single TSB plate containing either Cm or Kan. Once dried, plates were then incubated overnight or for two days depending on the strain being tested. While this protocol was being optimized, the plasmids pBBR122-CmR and pBBR122-KanR were used at 100 ng. Furthermore, while optimizing electroporation voltages, the OD of the final cell suspension was consistent across replicates in each strain but was not normalized to an OD of 25 for each strain.

### High-Throughput Expression Sequence Determination

Genomic DNA (gDNA) for each strain was extracted and verified using 16S PCR as described above. 90 µL of gDNA for each strain was then sonicated with a Covaris S220 ultrasonicator at a duty factor of 5 and 200 cycles per burst for 80 seconds. Fragmented gDNA was extracted at a size range of 100-600 bp from a 1% agarose gel run at 125 V for 20 minutes using Zymo’s gel extraction kit (D4001) according to the manufacturer’s protocol. The gDNA fragments were then end-repaired using the LGC Biosearch Technologies End-It DNA End-Repair Kit (ER81050) and subsequently purified using Zymo Clean and Concentrator-5 kit (D4004) according to the manufacturer’s instructions. The ES library vectors were prepared by inverse-PCR from the starting plasmids pBBR122-CmR-sfGFP or pBBR122-KanR-sfGFP. 1 μL of DpnI (R0176L) was added to the PCR reaction and incubated at 37 °C for 3 hours. After digestion, the backbone was purified using Zymo Clean and Concentrator-5 kit.

Ligation of the MRTK vector and gDNA fragments was performed using the LGC Biosearch Technologies Fast-Link DNA Ligation Kit (LK6201H) overnight at room temperature. The overnight ligation reaction was deactivated at 70 °C for 10 minutes and cleaned with Zymo Clean and Concentrator-5 kit. The ligation reaction was eluted twice with 6 µL of Nuclease Free water preheated to 55 °C. 5 µL of ligation reactions were transformed into their respective strains at their optimized voltages. 25 µL of the recovery solution was used to make serial dilutions to count transformants and the remaining volume was inoculated into 50 mL of TSB with either Cm or Kan and incubated overnight at 26 °C. The next day, any libraries in stationary phase were subinoculated at a ratio of 40:1 and incubated at 30 °C for 2 hours to reach log phase. Cells that were ready for sorting were harvested by centrifugation at 5,000 xg for 5 minutes and resuspended in 1x PBS.

Harvested ES libraries were sorted for green fluorescence using a BD FACSmelody. Sorting gates, voltages and thresholds were determined using the WT strain and strains expressing GFP as a positive control for each sorting experiment. Plasmid pBBR122-CmR-sfGFP was used as a positive control for sorting Elu cells, pBBR122-KanR-sfGFP’ was used as a positive control for sorting *Ppu* cells and pBBR122-CmR-sfGFP’ was used as a positive control for sorting *Hro*, *Bpi* and *Sma* cells. A minimum of 900 GFP positive events were sorted for each sorting run and then plated on TSB agar plates with either Cm or Kan (Table 1). Up to 96 fluorescent colonies were selected visually with a UV flashlight for additional screening and sequencing. Selected colonies were incubated in 2 mL TSB+Cm overnight at 30 °C and at 250 rpm. The next morning, the cells were sub-inoculated into 2 mL of fresh media at a 1:40 volume ratio and grown for 2-3 hours in the same conditions. The fluorescence of each culture was determined using a BD Accuri™ C6 Plus Flow Cytometer. A subset of these colonies with the largest range of fluorescence were then sent for sanger sequencing.

Plasmid DNA was extracted from cultures using Zymo’s Zyppy miniprep kit (D4020). DNA extraction followed the manufacturer’s protocol with the exception of adding 240 μL of the 7x Lysis buffer and 450 μL of Neutralization buffer. The promoter region of the plasmid was amplified using NEB Hot Start 2x Master Mix and sequenced via Sanger sequencing. Based on the Sanger sequencing results, ESs containing part of the downstream gene were repaired via the NEB Q5 mutagenesis kit according to manufacturer’s instructions. The resulting repaired ESs were then re-validated in biological triplicate in the same fashion as the initial selection of sorted sfGFP cells.

### Statistical analyses

All statistical analysis was performed using JMP Pro 17. One-way ANOVA analysis, Tukey Kramer, and HSU’s MCB were used to determine the best voltage and methylation pattern for each strain’s transformation protocol. Voltage optimization was performed with technical replicates of n = 5 or 6 where each replicate was a independent transformation from the same cell culture. The rest of the experiments were performed with biological replicates of n=3 where each replicate was a culture isolated from a single colony.

## Acknowledgements

All figures were created with Biorender and GraphPad. We are grateful to Dr. Ryan Paerl for training on the FACSmelody for expression sequence sorting experiments and Dr. Albert Keung for training and use of the Biorad FX384 Touch Real-Time PCR Detection System. We are also grateful to Dr. Justin Vento and Dr. Jennie Fagen from the Biesel lab for providing *Escherichia coli* 135 and the pBBR122 plasmid respectively, as well as Manuel Kleiner for providing the SynCom strains.

## Author Information

JVS, ZL and NC designed the experiments performed in this work. JVS performed all experiments, with the exception of qPCR which was performed by ZL. JVS and JC cloned the C plasmids of the MRTK. JC imaged the fluorescent SynCom strains.

## Ethics Declarations

The authors declare no conflicts of interest.

## SUPPLEMENTAL FIGURES

### Supplementary Excel file (Tables S1-S6)

**Figure S1:**
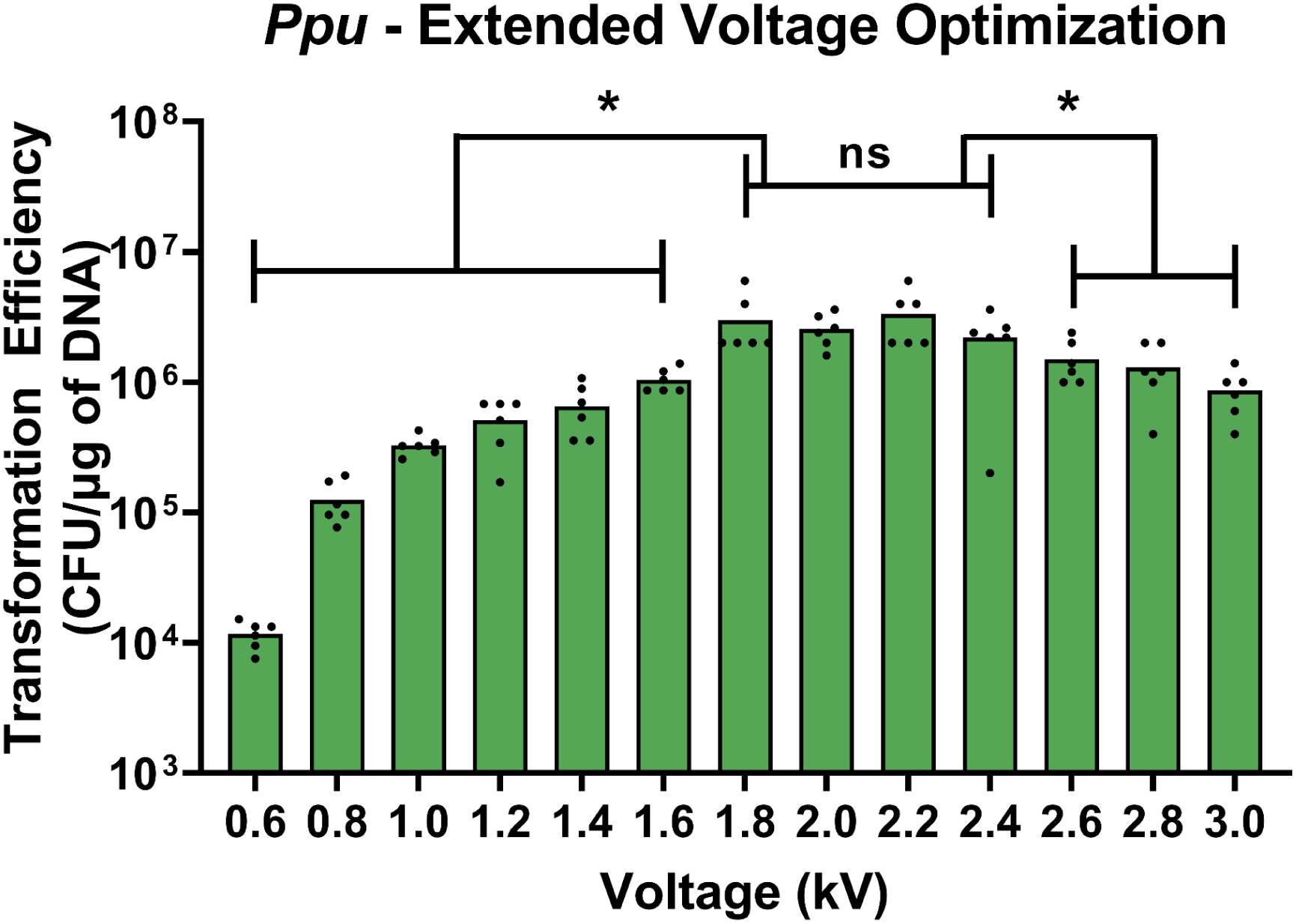
Extended Voltage Optimization for *Ppu.* Transformation efficiency of *Ppu* (CFU/μg of DNA) across an extended range of voltages. Statistical analysis was performed using One-way ANOVA with Tukey-Kramer post-hoc test to determine the optimal voltages. Data was collected in technical replicates of a size n=6.

**Figure S2:**
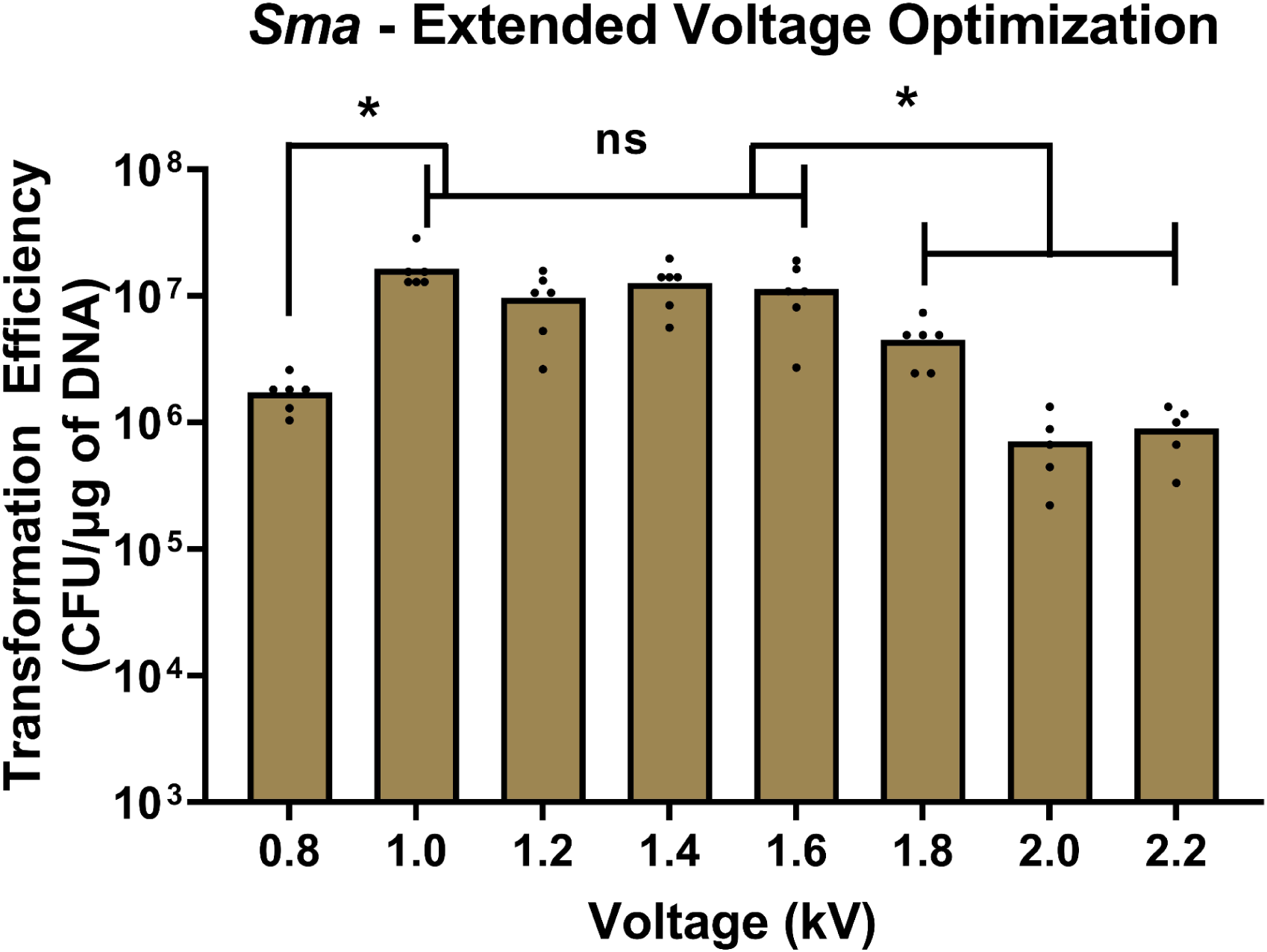
Extended Voltage Optimization for *Sma.* Transformation efficiency of *Sma* (CFU/μg of DNA) across an extended range of voltages. Statistical analysis was performed using One-way ANOVA with Tukey-Kramer post-hoc test to determine the optimal voltages. Data was collected in technical replicates of a size n=6.

**Figure S3:**
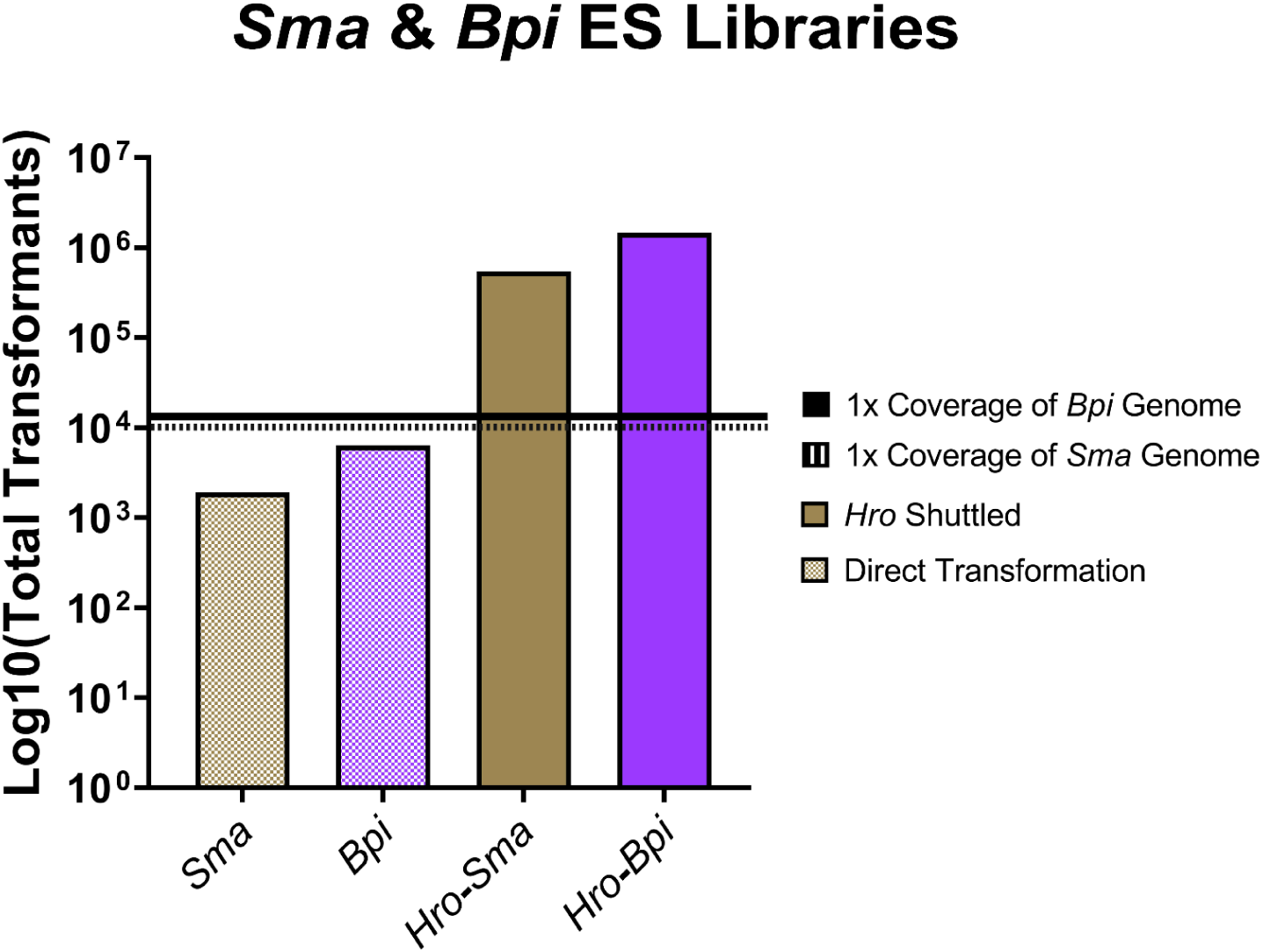
ES library sizes for Sma and Bpi. Total transformants of the ES libraries directly into *Sma* and *Bpi* and libraries shuttled through *Hro*. Directly transformed libraries have patterned bars and ES libraries shuttled through *Hro* are solid bars. 1x coverage of *Bpi*’s and *Sma*’s genome are shown by the solid and dotted lines respectively.

**Figure S4:**
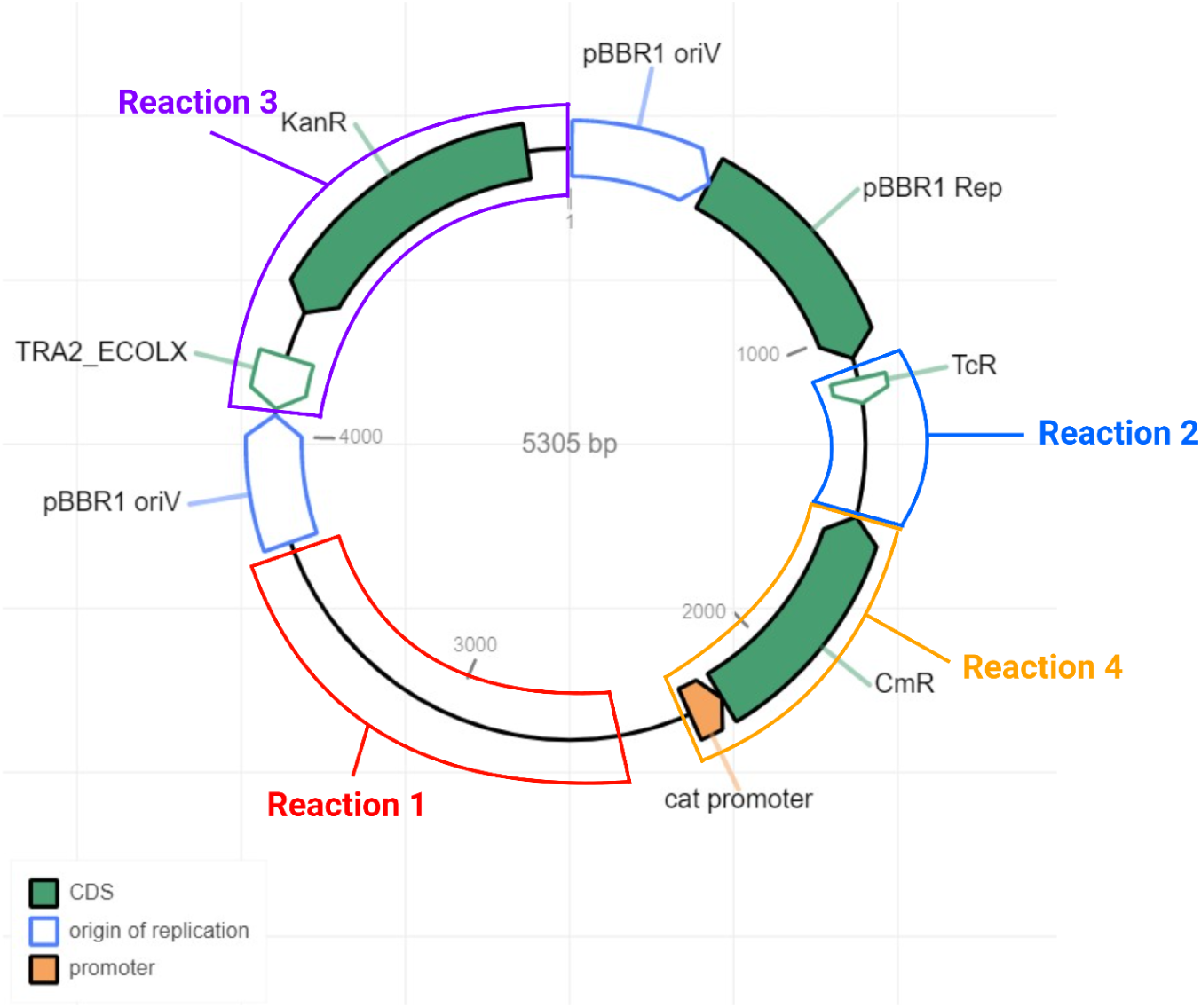
Deconstruction of pBBR122. A plannotate annotation of the plasmid pBBR122 with labeled segments of cloning reactions conducted to make the plasmids used in this study. Cloning reaction 1 removed the deactivated MobA gene, Cloning reaction 2 replaced this segment with 2 terminators, Cloning reaction 3 removed the Kanamycin resistance gene and Cloning reaction 4 replaced CmR with other selective markers.

